# A role for tectorial membrane mechanics in activating the cochlear amplifier

**DOI:** 10.1101/2020.05.06.080549

**Authors:** A. Nankali, Y. Wang, C. E. Strimbu, E. S. Olson, K. Grosh

## Abstract

The mechanical and electrical responses of the mammalian cochlea to acoustic stimuli are nonlinear and highly tuned in frequency. This is due to the electromechanical properties of cochlear outer hair cells (OHCs). At each location along the cochlear spiral, the OHCs mediate an active process in which the sensory tissue motion is enhanced at frequencies close to the most sensitive frequency (called the characteristic frequency CF). Previous experimental results showing an approximate 0.3 cycle phase shift in the OHC-generated extracellular voltage relative the basilar membrane displacement that is initiated at a frequency approximately one-half octave lower than the CF are repeated in the present paper with similar findings. This shift is significant because it brings the phase of the OHC-derived electromotile force near to that of the basilar membrane velocity at frequencies above the shift, thereby enabling the transfer of electrical to mechanical power at the basilar membrane. In order to seek a candidate physical mechanism for this phenomenon, we used a comprehensive electromechanical mathematical model of the cochlear response to sound. The model predicts the phase shift in the extracellular voltage referenced to the basilar membrane at a frequency approximately one-half octave below CF, in accordance with the experimental data. In the model, this feature arises from a minimum in the radial impedance of the tectorial membrane and its limbal attachment. These experimental and theoretical results are consistent with the hypothesis that a tectorial membrane resonance introduces the correct phasing between mechanical and electrical responses for power generation, effectively turning on the cochlear amplifier.

**SIGNIFICANCE:** The mechanical and electrical responses of the mammalian cochlea are nonlinear exhibiting up to a thousand-fold difference in gain depending on the frequency and level of sound stimulus. Cochlear outer hair cells (OHC) are broadband electro-mechanical energy converters that mediate this nonlinear active process. However, the mechanism by which the OHC electromotile force acquires the appropriate phase to power this nonlinearity remains unknown. By analyzing new and existing experimental data and using a mathematical model, we address this open issue. We present evidence which suggests that a relatively simple feature, the frequency dependence of the radial impedance of the tectorial membrane, provides requisite mechanics to turn on the frequency-specific nonlinear process essential for healthy hearing.

## INTRODUCTION

### Background

The pressure difference across the sensory tissue of the cochlea, the organ of Corti complex (OOC, Fig. 1), produces vibrations that ultimately give rise to the sensation of sound. The OOC motions are boosted by a nonlinear active process that enables sound processing over a broad range of frequencies and intensities (1). Somatic motility of the mechanosensory outer hair cell (OHC) is largely accepted as the key mediator of the active cochlear mechanism (1–3). The electromechanical properties of the OHCs convert electrical energy, stored within a metabolically–maintained resting potential inside the cochlea, into mechanical energy. The active process causes a nonlinear response such that BM gain relative to the stapes motion is on the order of 10^4^ at low sound pressure levels (SPLs), while at high SPLs the gain is on the order of 100 (4–6). This nonlinearity compresses the dynamic range by two orders of magnitude, maintaining sub-nanometer sensitivity for threshold–level sounds while protecting the delicate sensory microstructures at high sound pressure levels. The onset of nonlinearity in BM responses measured at mid-to-high characteristic frequency (CF) locations occurs at frequencies about ½ octave below the CF and extends to slightly above CF in most rodents (4, 6–9). Experiments in living cochleae, (6, 10–13) and *in vitro* (14) have demonstrated that OHCs generate forces over a wide frequency range. The nonlinear, frequency-location-specific BM-motion enhancement, sometimes termed the cochlear amplifier, is studied in the current paper.

**Figure 1:**
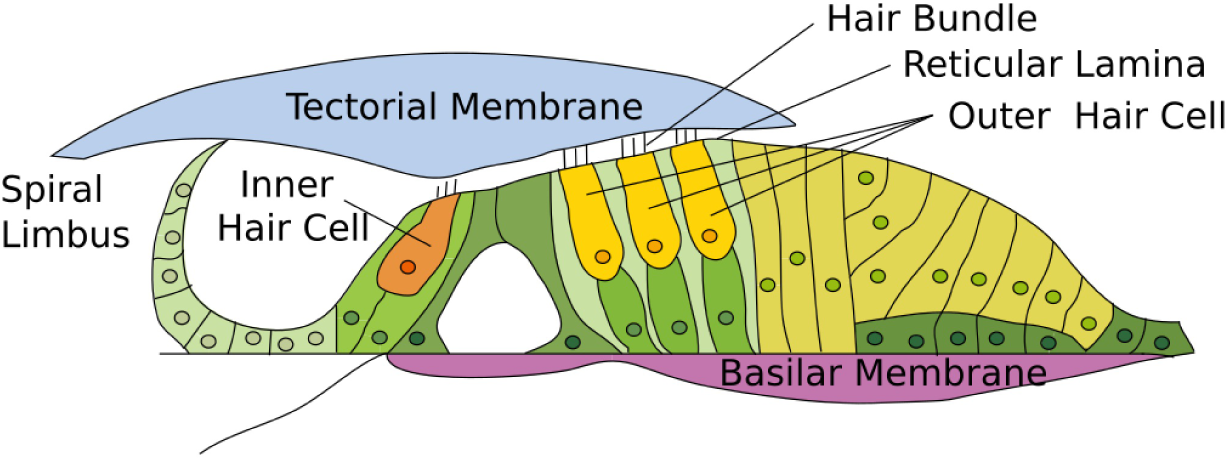
organ of Corti complex (OCC), refers to the cellular organ of Corti and the acellular tectorial and basilar membranes (TM and BM, respectively).

The main motivation for the present work stems from Dong and Olson (5), who explored cochlear amplification by measuring sound-evoked electrical and mechanical responses *in vivo*. A specialized dual-pressure-voltage sensor was used to measure the scala tympani (ST) voltage and acoustic pressure (*P*_*st*_) simultaneously at the same location close to the BM. Further, pressure differences were used to make an approximate measurement of BM displacement. We will denote the ST voltage measured close to the BM as the local cochlear microphonic (LCM); this potential closely tracks the transducer current flowing through OHCs in the vicinity of the electrode (15, 16).

Fig. 2*A* and *B* revisits frequency response results from the Dong and Olson study (experiment wg165) (5). In this figure, the LCM evoked by different SPL (thin lines) is compared to a pressure-based estimate of the BM displacement (thick lines). Because of the approximate nature of the pressure-based displacement measurement, we also compare the LCM to BM displacement data acquired in a different manner in Fig. 2*C* and *D*. Here,the LCM from wg165 is compared to a separate BM displacement data set from the same lab, but from a different animal (GB800), gathered using a relatively straightforward laser-based interferometric method (but without simultaneous LCM) as described in (12) and briefly in Methods section. The measurement location for both experiments is similar. The data are shown on a frequency axis that is normalized to the CF of each preparation, 23 kHz for wg165 and 26 kHz for GB800.

**Figure 2:**
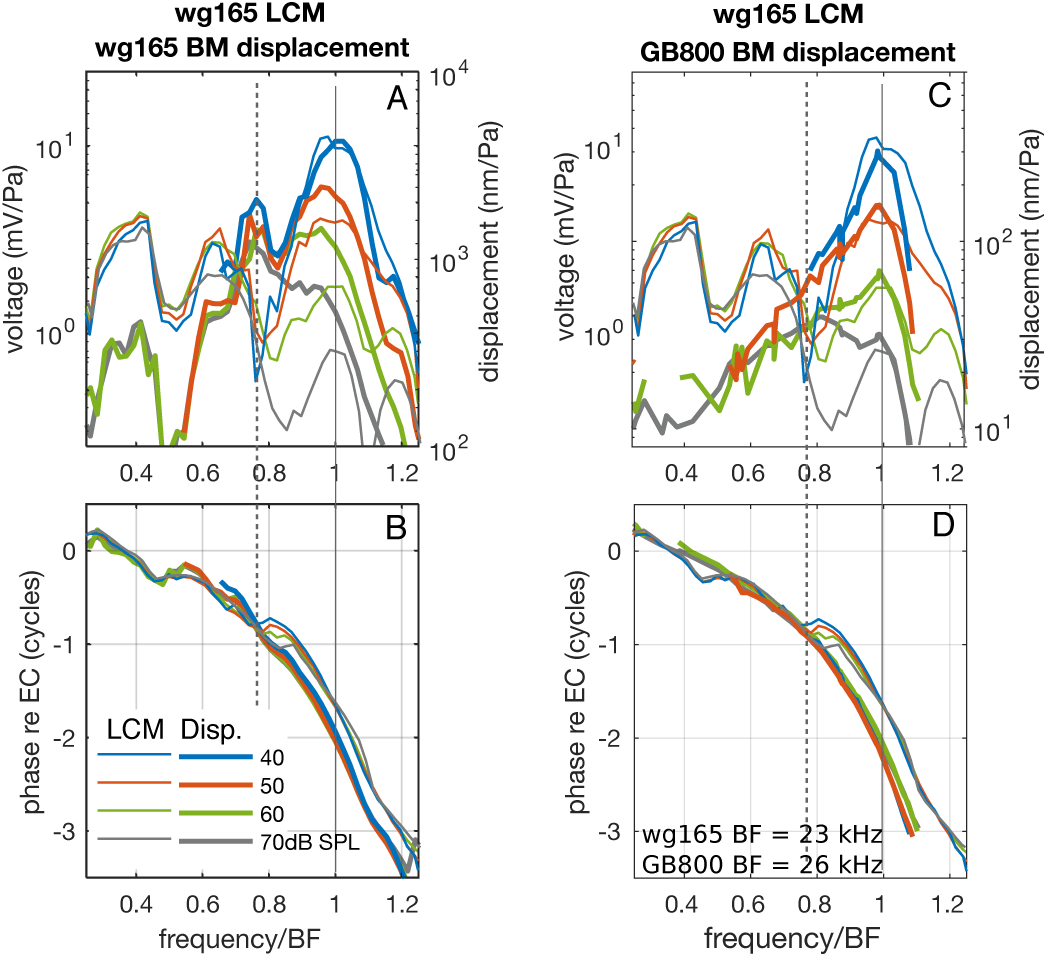
(A and B) Previously published data (5) (exp.wg165) in which LCM and displacement were measured using a dual pressure-voltage sensor. All quantities are referenced to EC pressure. (A) Magnitude of LCM (thin) and BM displacement (thick). (B) Phase of LCM (thin) and BM displacement (thick). (C and D) Here LCM from wg165 are plotted with displacement found with a laser-based method (exp.GB800). These panels are included to reinforce the findings of A and B. (C) Magnitude of LCM (thin) and BM displacement (thick). (D) Phase of LCM (thin) and BM displacement (thick). Positive displacement direction is up in Fig. 1, in the direction of the scala media.

The aspect of the data that is of primary interest is the phase of the LCM relative to BM displacement in Figs. 2B and *D*. The LCM phase rides along with the displacement phase up to a frequency of ∼ 0.76 *f*_*CF*_, but then shifts upward to lead the BM phase. At a frequency ∼ 0.9 *f*_*CF*_, the lead of LCM re BM displacement is ∼ 0.27 cycle for the simultaneous wg165 LCM–pressure data of Fig. 2*B*, and ∼ 0.4 cycle for the data of Fig. 2*D*, in which LCM data from wg165 are compared to BM displacement data from GB800. Comparisons of the LCM phase to either the simultaneously measured wg165 BM displacement or the displacement phase data from a different animal (GB800) both show that the displacement and LCM phase track closely up to the bifurcation point at ∼ 0.76 *f*_*CF*_. This consistency confirms that this finding is not related to the non-standard displacement measurement in experiment wg165. Moreover, this correspondence shows that we can use the phase information from one gerbil’s BM displacement as a reference for the LCM of a different animal, provided the locations are nearby as we have used in Fig. 2. The second interesting aspect of the data is that the magnitude of the LCM exhibits a notch corresponding to the phase shift at ∼ 0.76 *f*_*CF*_. This is clear at SPLs up to 60 dB SPL, but less so at 70 dB SPL. Hence this effect seems to saturate at high SPL in the experiments.

The significance of the phase shift was analyzed in (5), and shown to correspond to a transition from non-effective forcing by OHC somatic forces below the shift frequency (here 0.76 *f*_*CF*_) to effective forcing and power generation in the CF region. This analysis used previously-published relationships between OHC somatic force and OHC voltage (14) and between LCM (representing OHC current) and OHC voltage (17). The conclusion that above the transition frequency OHC forcing becomes effective on the BM is consistent with the observation that the BM gain curves bifurcated from their sub-CF linear backbone at ∼ 0.76 *f*_*CF*_. Thus, cochlear amplification, defined as the nonlinear peaking of the BM gain, began at the frequency of the LCM amplitude notch and phase shift relative to BM motion; these two transition frequencies coincided. The existence of a transition frequency that marks the onset of nonlinearity is commonly seen in the BM response to acoustic stimulation. It is seen for example, in gerbils (7), guinea pigs (8, 9) and mice (6) (see Table 1).

**Table 1:** Comparison frequency of the onset of nonlinearity (*f*_*NL*_) to the CF of the measurement location from different experimental data

| Animal | CF(kHz) | $f_{NL}$(kHz) | $f_{NL}/CF$ | Source |
| --- | --- | --- | --- | --- |
| Chinchilla | 10 | 7 | 0.7 | (4) |
| Gerbil | 23 | $\sim 17$ | 0.7 | (5) |
| Mouse | 10 | 6 | 0.6 | (6) |
| Guinea pig | $\sim 15$ | $\sim 10$ | 0.7 | (9) |

The physical basis for the amplifier-activating phase shift is the subject of the current paper. We present simulations from a realistic cochlear model to explore the basis for the experimental findings. Many models of the cochlea have been developed. They include lumped element circuit models with and without active elements (18–20), and detailed FEM models (21–24). Our model couples a 3D FEM-based treatment of the fluid with a circuit model of the sensory tissue. The key feature of our model is that it is fully electromechanically coupled. Mechanical motion activates the mechanoelectric transduction in the OHC hair bundles which gives rise to the current that drives OHC basolateral electromechanical forcing. The amplitude and phase of the OHC activity are not *a priori* assumed, but rather are an output of the model. Hence, this model enables us to seek a causal relation (inside the model) between output responses and mechanisms. In addition to showing model results, we reinforce the experimental finding by showing new LCM data that confirm the findings of phase shift and amplitude notch, and extend the previous findings into a lower frequency region. Additional supportive experimental evidence has already been published (12, 25, 26). All of these results are from gerbil. A similar LCM amplitude notch and phase shift was observed in 1976 work by Yates and Johnstone in guinea pig (27) but was not apparent in the more recent work of (16). Further studies are needed to determine the species generality of the findings.

## MATERIAL AND METHODS

### Experimental measurements of local cochlear microphonic and BM motion

This paper is primarily a modeling paper, with experimental data included to bolster previous experimental findings. The wg165 data of Fig. 2 and the BM motion data of Fig.4 (28) were previously published. Other data from these figures are unpublished, although similar LCM and OCT-based displacement data have been presented and methods fully described in recent work (12, 29). To keep the focus on the modeling results, the description of experimental methods for the unpublished data is kept short. Procedures were approved by the Columbia University Institutional Animal Care and Use Committee. Young adult gerbils were sedated with ketamine and anesthetized with Pentobarbital, with supplemental dosing throughout the experiment and the analgesic Buprenorphine was given every six hours. Animals were euthanized at the end of the experiment. The stimulus generation and acquisition were performed using MATLAB-based programs and a Tucker Davis Technologies (TDT) System. Sound stimulation was generated via an electrically shielded Fostex dynamic speaker, connected in a closed-field configuration to the ear canal (EC). The sound calibration was performed within the EC using a Sokolich ultrasonic probe microphone. Pure tone stimuli were used for LCM measurements and multi-tone stimuli were used for the BM motion measurement of Fig.2. The phase response that is the focus of the current paper is not significantly affected by multi-tone versus single-tone stimulation (12, 30).

#### Local cochlear microphonic

For LCM measurements, after opening the bulla, a hole of diameter ∼100 *µ*m was hand-drilled to access ST through the bony wall of the first turn of the gerbil cochlea where the characteristic frequency (CF) was 15 - 25 kHz. A polymer-coated tungsten electrode (FHC Inc., Bowdoin Maine) with shank diameter 250 *µ*m and tip diameter ∼1 *µ*m, held in a motorized micromanipulator (Marzhauser) was inserted into the hole and used to measure voltage responses to acoustic stimuli. (In (5) the voltage sensor was an insulated wire electrode adhered to the side of a pressure sensor.) The impedance of the electrode was 1 - 5 M Ω at 500 Hz. The metal electrode had a broad-band frequency response, and thus no correction due to low-pass filtering by the electrode was needed (5, 31). The voltage was amplified × 500 or 1000 by a PARC *EG*&*G* amplifier. A reference electrode was placed on the muscle at the neck. Once within the cochlea the electrode was advanced in steps toward the BM and the responses to acoustic stimulation measured. When traveling wave responses were detected through several cycles, the measurement was deemed “local”.

#### BM motion

A commercial ThorLabs Telesto III spectral domain optical coherence tomography (SD-OCT) system was used to measure the vibrations of the BM through the intact round window membrane. OCT-based measurements are a laser-based motion measurement system and the measured displacement is based on the strict physical quantity of light wavelength. In Fig.2 *C* and *D* we included one data set taken with the Telesto, to support the results of Fig.2 *A* and *B*, in which the displacement was measured with a less stringent method, via fluid pressure differences. Data acquisition and analysis scripts were written in Matlab (R2016b) and in C++, based on the Thorlabs Software Development Kit. To make a measurement with the Telesto, first a two-dimensional scan, termed a B-scan, was taken across the radial direction of the OCC to image a radial section of the BM and OC (as in the cartoon of Fig.1). Then scanning was arrested so that the OCT collected data along one axial line, termed an A-scan. While acoustically stimulating the ear with a multi-tone stimulus, the OCT system acquired a series of time-locked A-scans, termed an M-scan, at a sample rate of ∼ 100 kHz. After initial processing of the M-scan, selected pixels in the A-scan can be chosen for extraction of the displacement vs. time. Locations along the A-scan include structures within the OC, but for the purposes of the the present paper, the motion at an A-scan pixel corresponding to the BM was shown in Fig 2.

#### Combining LCM and BM motion from different experiments

In Fig. 2 *C* and *D* and Fig. 4, LCM and BM motion responses responses from different animals are compared. In order to make the comparison, measurements with reasonably similar CFs were paired and the frequency axis was normalized by CF. A BM motion data set with BF of 26 kHz was used for Fig. 2 *C* and *D* and a single BM motion data set, with BF of 15.5 kHz was used for all the comparisons in Fig. 4. The *f* /CF normalization is the only manipulation required for Fig. 2 *C* and *D*. For the results of Fig. 4, the approximately CF-matched results came from BM motion presented in (28). In that data set BM velocity was measured relative to stapes, and a 25 *µ*s middle ear delay (32) and 0.25 phase shift is applied to convert the data to displacement relative to EC pressure.

**Figure 4:**
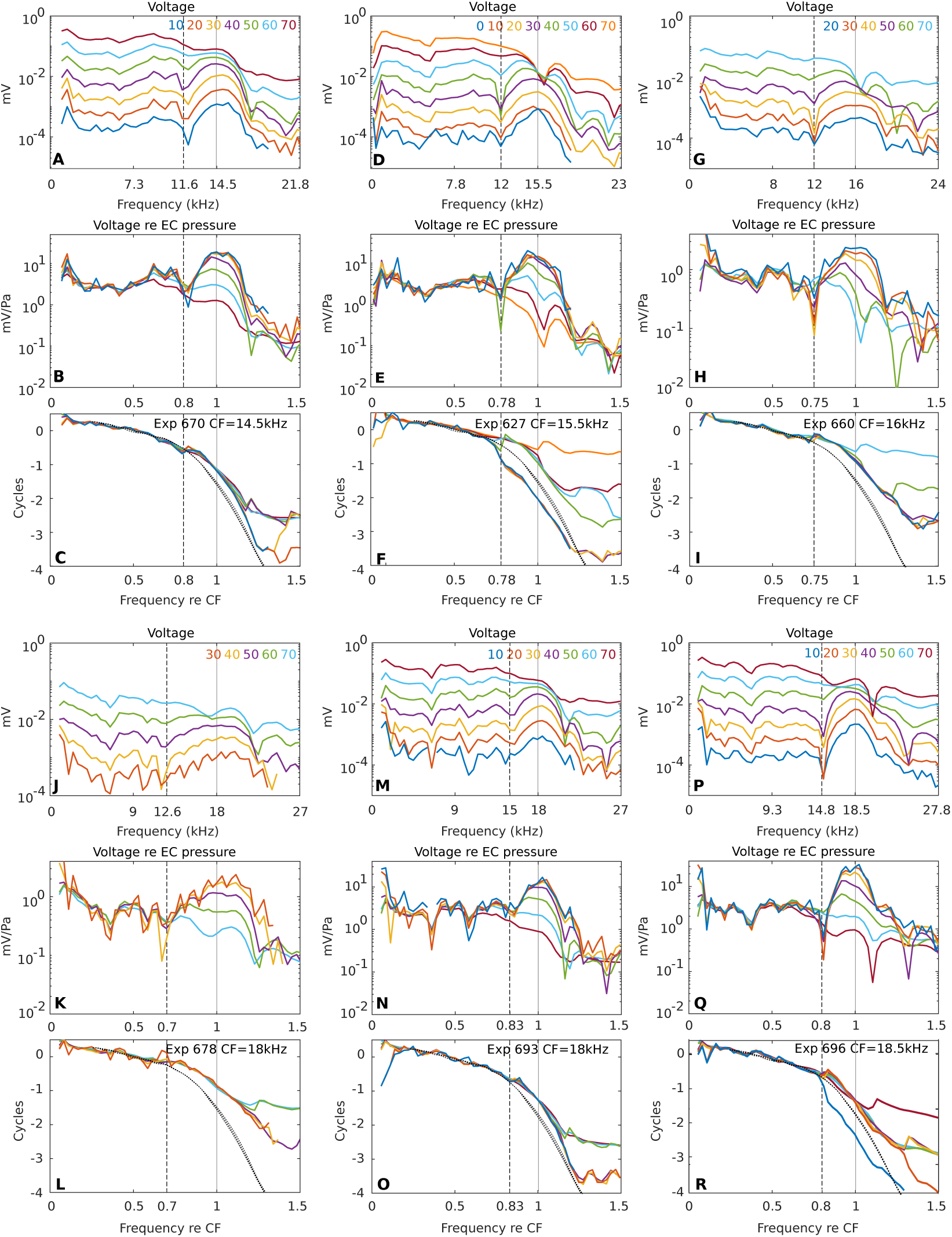
Frequency response of LCM in six preparations. Each preparation is represented in three vertical panels. The color key to SPL is in the top panel, the CF and animal number is noted in the bottom panel. (A,D,G,J,M,P) Response magnitude. Horizontal dashed line is the noise floor. (B,E,H,K,N,Q) Magnitude normalized to EC pressure. (C,F,I,L,O,R) Phase referenced to EC pressure. In (B,E,H,K,N,Q,C,F,I,L,O,R) the frequency axis is normalized to CF. To provide a comparison similar to that of Fig.2, in panels C,F,I,L,O and R phase data from BM motion responses from (28) are included in thin dashed black lines. The CF of the BM motion data was 15.5 kHz. 30, 40 and 50 dB SPL results are included to indicate SPL was not critical to the comparison.

### Mathematical Model

A physiologically-based 3D model of the cochlea was used to probe the basis for the experimental observations of LCM and BM motion. The model was originally designed to analyze guinea pig data (33, 34). Variations of the model have been used to probe responses in mice with genetically manipulated limbal attachments of the TM (35); the model predicted that a mammal with a TM detached from the limbus but still connected to the OHC stereocilia would possess active amplification and nonlinearity with sharper tuning than the wild-type animal, as had been found experimentally (36). In recent works, the model has been used to explore both apical and basal cochlear responses (23) and responses in *in vitro* preparations (24). Hence, this model can be considered a general model of the mammalian cochlea suitable for studying mechanisms of function.

We recount the main features of the model next (33, 34). Fig. 3 shows a schematic of the cochlear box model (panel *A*) along with the OCC components (panel *B* and *C*). In Fig. 3A, the macroscopic fluid-structure configuration is shown. The scala vestibuli (SV) and scala media (SM) fluids are taken together for fluid-mechanical purposes. Electrical cables are present in each of the fluid scala (SV, SM, and ST) and are used to model the ionic current flow in each (see Fig.S1A in the Supplemental Information). Fig. 3*B* shows a cross–sectional view of the OCC and surrounding cochlear fluids. The difference in the ST and SV pressure across the BM causes the OCC to vibrate in response to excitation at the stapes. Structural longitudinal coupling is included in the BM and TM mechanics (34). In the model used in the present paper, the fluid is modeled as inviscid and incompressible (34, 37), except in the subtectorial space. There, viscosity is incorporated through fluid shearing between the TM and the reticular lamina (RL). In addition, a small amount of structural damping of the BM and TM are also included. As schematically shown in Fig. 3*B*, the microstructural components of the OCC are coupled to each other. The TM is anchored to the spiral limbus via a spring with stiffness *k*_*tms*_ and connected to the hair bundles (HB) of the OHCs through a stiffness *k*_*hb*_ (shown as a spiral spring at the base of the HB in Fig. 3B). The three primary dependent structural variables shown in Fig. 3*B* are the displacement of the BM (*u*_*bm*_), the shear displacement (*u*_*tms*_) and the transverse displacement (*u*_*tmb*_) of the TM. Other structural displacements are then computed through kinematic constraints (33).

**Figure 3:**
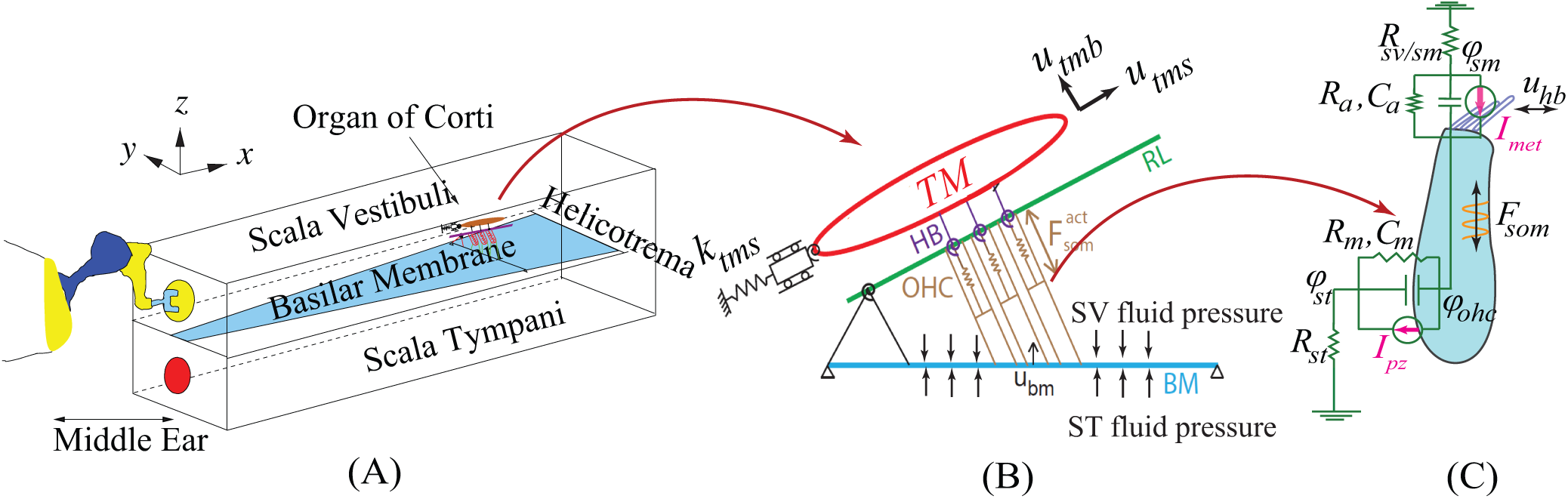
(A) The cochlear box model. For visual clarity the organ of Corti is only pictured at one cross section. (B) A schematic of the transverse section of the OCC microstructure. The hair bundles of the OHCs are shown connecting the RL to the TM through springs (spring constant *k*_*hb*_). The TM is anchored to the spiral limbus via dashpot and spring *k*_*tms*_. (C) Schematic of an OHC and the mechano-electric-transducer (MET) apparatus. The OCC cross-sections are coupled mechanically through longitudinal coupling in the TM and BM, the 3D-fluid pressure, and through electrical coupling (the three cable model models in the scala). (abbreviations: TM, tectorial membrane; OHC, outer hair cell; RL, reticular lamina; BM, basilar membrane).

Fig. 3*C* shows the local electrical circuitry of the OHC and the coupling of the tip displacement the hair bundle (HB) relative to the RL and somatic strain to the electrical domain. The deflection of the HBs triggers the opening of the MET channels resulting in current flow, *I*_*hb*_, into the OHC. This nonlinear process has been linearized as in (38):

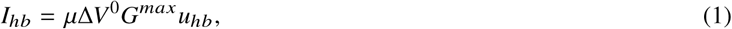

where *G*^*max*^ is the maximum saturating conductance of the HB and Δ*V* ^0^ is the resting value of the voltage difference between scala media (SM) and intracellular OHC potential. The MET scaling factor, *µ*, controls the sensitivity of the MET channels; it varies from 0, representing a nearly passive model, to 1, fully active (on the stability boundary). The model is quasi-linear in that varying *µ* simulates the SPL-dependent saturating nonlinearity of the MET channels. This quasi-linear approximation has been shown to be a good approximation with pure tone acoustic input (38). The OHC HB tip displacement relative to the RL (*u*_*hb*_) is a linear combination of the BM transverse (*u*_*bm*_), TM shear (*u*_*tms*_) and TM bending (*u*_*tmb*_) displacements (see (33) and Fig. 3.)

The coupled electromechanical relations governing the total axial compressive force (*F*_*ohc*_) and transmembrane current (*I*_*ohc*_) of the OHC are given by

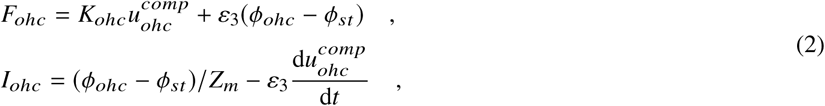

where 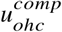 is the OHC compression (the difference between the displacement of the apical and basal poles of the OHC), (*ϕ*_*ohc*_ – *ϕ*_*st*_) is the transmembrane potential (the difference between the intracellular potential *ϕ*_*ohc*_ and extracellular potential *ϕ*_*st*_). The transduction current, *I*_*hb*_, activates the OHC somatic motility, which applies a mechanical force on the BM and RL. The OHC somatic force, *F*_*som*_, is proportional to the product of the OHC transmembrane potential and the piezoelectric coupling coefficient, *ε*_3_: *F*_*som*_ = *ε*_3_(*ϕ*_*ohc*_ − *ϕ*_*st*_). In Eq. 2, *Z*_*m*_ is the OHC basolateral electrical impedance (see Fig. 3) and *K*_*ohc*_ represents the OHC stiffness. 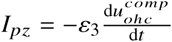 corresponds to the total current due to the piezoelectric-like behavior of the OHC. Hence, the mechanical degrees of freedom give rise to electrical currents (i.e., *I*_*hb*_ and *I*_*pz*_ are due to mechanical motion) and electrical degrees of freedom give rise to mechanical forcing (*F*_*som*_). The mathematical model outlined above gives rise to a set of coupled partial differential equations, which are solved using the finite element method (39). The parameters of the model follow (35) with the exception of those listed in Table 2. We also study the impact of manipulating the damping coefficients in this paper

**Table 2:** Parameter values that are changed from (35). *x* is distance from stapes (meters).

| Property | Description | Value |
| --- | --- | --- |
| $K_{tms}$ | TM shear stiffness per unit length | $3 \times 10^4 e^{-3.75x} \text{ N/m}^2$ |
| $C_{tms}$ | TM shearing damping coefficient per unit length | $0.006 \text{ N s/m}^2$ |
| $C_{tmb}$ | TM bending damping coefficient per unit length | $0.04 \text{ N s/m}^2$ |
| $C_{bm}$ | BM damping coefficient per unit length | $0.05 \text{ N s/m}^2$ |

## RESULTS

### Experimental LCM and BM displacement phase comparison

Fig. 4 shows the frequency response of the tone-evoked LCM over a range of SPLs for six preparations, with the results from each preparation comprising three vertically-stacked panels. For example, Fig. 4 *A-C* are from expt. 670, with *A* the unnormalized amplitudes, *B* the amplitudes normalized to the EC pressure and *C* the phase relative to the EC pressure. In *B* and *C* the abscissa is normalized to the peak frequency of the lowest SPL response, the CF. The low and moderate SPL responses show several cycles of traveling wave phase accumulation, indicating that the voltage responses were dominated by local OHCs (5). The CFs range from 14.5 to 18.5 kHz. In the phase panels, we also present the BM displacement phase responses from (28) at three different SPLs as thin dashed black lines; this phase is seen not to vary significantly with the SPL. The same BM displacement phase is shown in each phase panel, and this enables a common reference for the comparison LCM phase to the BM displacement phase similar to that of Fig. 2. The CF of the BM displacement data was 15.5 kHz, close to the CF of the LCM measurements.

The features we are most interested in are the magnitude notch (local minimum) in LCM and the concomitant shift of the phase difference between LCM and BM displacement. The magnitude notch is clear in five of the seven LCM experiments presented in this paper (Expts. 165, 627, 660, 693, 696) and mild in two others (Expts. 678 and 693). The location of the amplitude notch does not change with SPL but its depth varies and sometimes disappears at higher SPL values, as noted when describing Fig. 2. At the same frequency as the LCM notch, a shift of the LCM phase with respect to the BM displacement phase is apparent in all the experiments. We denote this shift frequency, *f*_*shift*_, using a dashed vertical line in the panels of Fig. 4. For some experimental results and specific SPL values, the phase shift is large and an unwrapping ambiguity can occur close to *f*_*shift*_ (as in panels F and R). At frequencies above these ambiguities, the phases at different SPL become offset by approximately a full cycle.

Including data from Fig. 2 and that from Fig. 4, we have an N of 8 to consider in order to extract two metrics from these data. (I) The phase transition occurs at a frequency relative to CF of 0.77 ± .04 (mean ± standard deviation). (II) Evaluated at 0.9 *f*_*CF*_, the phase lead of LCM relative to BM displacement is 0.37 ± .09 cycles. The transition frequency, *f*_*shift*_, divides the (nearly) linear sub-CF region and the peaked and nonlinear CF region. At SPLs 70 dB and above, nonlinearity began to extend into the sub-CF region, likely due to saturation of OHC current at relatively high SPL. As an aside, recent measurements of motions within the OCC in the base of the gerbil cochlea show that nonlinearity exists in the motions of intra-OCC structures at sub-CF frequencies (10–12). The sub-CF nonlinearity observed in the motion is similar to our observations of high SPL nonlinearity in sub-CF LCM (12), and indicates that the OHC electromotile force is present for all frequencies, but amplifies BM motion only in the peak region.

To summarize, the experimental results of Figs. 2 and 4 reinforce the findings of (5) in showing that at the frequency where the BM nonlinearity begins, a phase shift of OHC voltage relative to BM displacement occurs, “activating” the cochlear amplifier. In the simulations below we show how this activation of the cochlear amplifier could occur.

### Simulations

To interpret the experimental results, we created an analogous set of predictions using the cochlear model described in the Methods section. In Fig. 3*B*, a schematic of the cochlear cross section is shown along with the definitions of the directions of positive displacement for the BM (*u*_*bm*_) and the TM in the shear (*u*_*tms*_) and bending (*u*_*tmb*_) directions. We also compute the hair bundle deflection (*u*_*hb*_) that divided by the HB height equals the rotation of the HB relative to the cuticular plate (see Fig. 3*C*); this motion gives rise to the MET currents (*I*_*hb*_ in Eq. 1), and is a linear combination of *u*_*bm*_, *u*_*tmb*_, and *u*_*tms*_ (33). The theoretical prediction of the LCM is written as *ϕ*_*st*_. *ϕ*_*st*_ is strongly correlated to OHC current, thus the amplitude and phase of *ϕ*_*st*_ is expected to be similar to *u*_*hb*_. Deviations from similarity arise due to current spread from adjacent locations, which is included via the electrical cable model of the fluid spaces. The effect of current spread was studied previously (5), and is explored further in the Supplemental Information of the present paper. In Fig. 5, predictions of the ST voltage (*ϕ*_*st*_, see Fig. 3*C*) and mechanical responses (*u*_*bm*_, *u*_*hb*_, and *u*_*tms*_) to acoustical stimulation are shown for a basal region of the model (4 mm from the stapes). Frequencies in this plot are normalized to the BM peak frequency at low SPLs (CF), and amplitudes are presented relative to the stapes displacement. As in the experiment, a local minimum is seen in *ϕ*_*st*_ at a frequency below the CF, 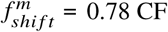 CF (the model-shift frequency), and the BM response does not evince a notch (consistent with the experimental results). From the phase plot, it can be seen that the phase of *ϕ*_*st*_ and *u*_*hb*_ deviate from *u*_*bm*_ at a frequency slightly less than that of the notch. Hence, the phenomenon of a notch and phase-shift frequency is predicted by the model.

**Figure 5:**
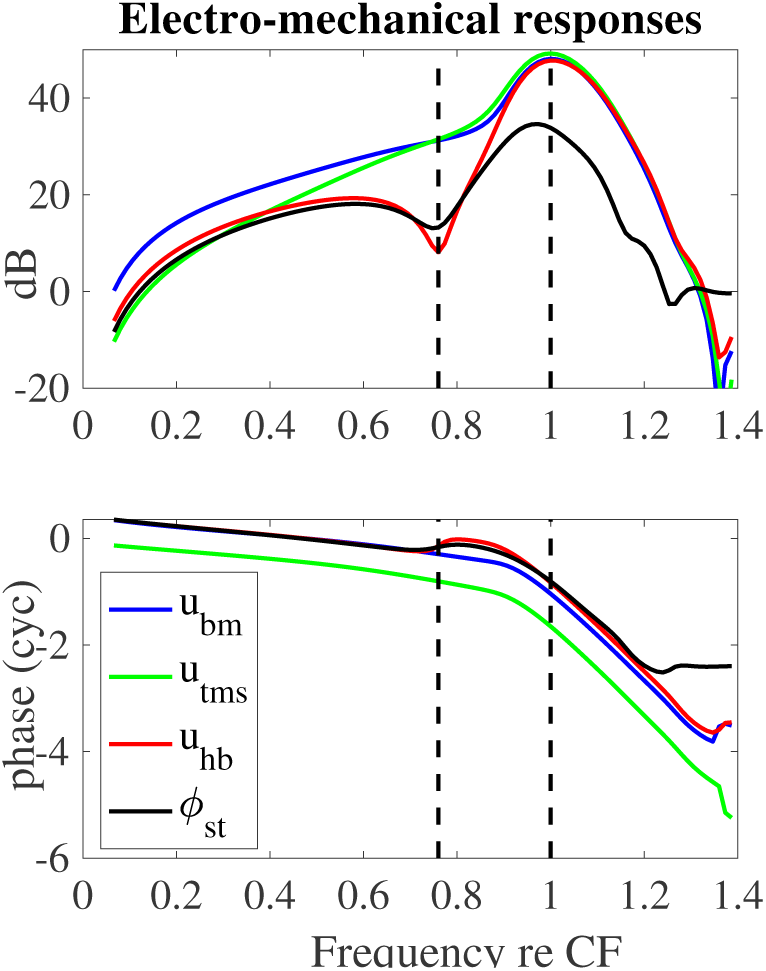
Model predictions of the transfer function for ST voltage (*ϕ*_*st*_), BM transverse (*u*_*bm*_), TM shear (*u*_*tms*_) and HB deflection (*u*_*hb*_) displacements relative to the stapes displacement; frequencies are normalized to the CF=14.8 kHz, the MET scaling factor is set to *µ* = 0.7, and the stapes positive motion convention is outward. The units for the ST voltage to stapes motion gain is in mV/nm in order to scale with the non-dimensional displacement gains.

To determine if these relationships are level dependent, the model predictions of the magnitude and phase of *ϕ*_*st*_ and *u*_*bm*_ gain (relative to the stapes displacement) are shown in Fig. 6 for a range of input SPL. Increasing SPL was simulated in our model by decreasing the MET sensitivity as embodied by the scaling factor, *µ* (see Eq. 1). Fig. 6*A* shows that the model-predicted voltage notch is present for all SPLs, unlike the experimental result where the notch is washed out for stimulus levels above ∼ 60 dB SPL. Hence, there are stimulus level variations in the cochlea that are not encompassed by simply varying *µ*. As in the experimental LCM results, the *ϕ*_*st*_ gain decreases with increasing SPL. At low frequencies nonlinearity is more pronounced in the model than in these single-tone experimental data, but in experiments with multi-tone stimuli (12), low frequency nonlinearity is strong. Hence, this difference between experiments and model results is considered a fairly minor quantitative difference. Strong nonlinearity is seen over all stimulus levels near CF in the model, as in the experiments. Fig. 6*B* shows amplitude and phase of the BM displacement as a function of SPL and frequency. As in the experiment (see Fig. 2*B*), the BM nonlinearity emerges strongly at a frequency near the notch frequency.

**Figure 6:**
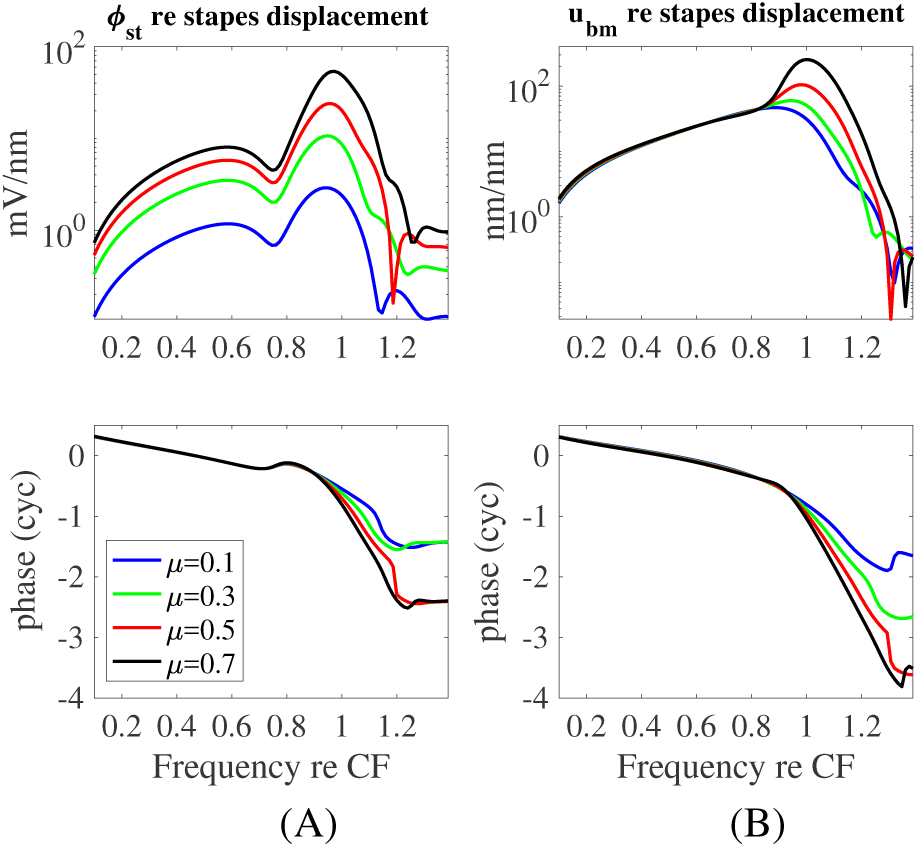
Theoretical predictions of the magnitude and phase of (A) the amplitude of the transfer function between *ϕ*_*st*_ and the stapes motion and (B) the BM displacement gain (relative to the stapes displacement) as a function of SPL and frequency (normalized to the CF=15 kHz). Variation of the input SPL on the gain is modeled by altering the MET sensitivity controlled by the parameter *µ* defined in Eq. 1. The voltage responses show the notch at 0.76 CF kHz while the onset of the BM nonlinearity occurs at slightly higher frequencies close to 0.82 CF (where low *µ* corresponds to a higher SPL as described in the Materials and Methods). The frequency of the notch is relatively independent of SPL. This is consistent with the measurements up to 70 dB SPL in Fig. 4 and 85 dB in Fig. 2*B*

In Fig. 7 we plot the measured (from (5), redrawn in our Fig. 2) and predicted phase difference between *ϕ*_*st*_ and *u*_*bm*_. The measured phase difference (dashed line in Fig. 7) underwent a phase shift at the transition frequency (0.76 CF) that produced a ∼ 0.34 cycle shift, when measured at 0.9 CF. The phase shift results (transition frequency and phase shift magnitude) from Fig. 4 were similar, as reported above. The theoretical prediction showed a phase shift with very similar onset and slope to the experimental value. The value of the phase shift at 0.9 CF was of ∼ 0.25 cycle, slightly smaller than the experimental value of 0.37 ±0.9 cycle. Above the CF the phase difference in the both experimental and modeling responses underwent more extreme variations that are associated with the onset of the supra-CF phase plateau.

**Figure 7:**
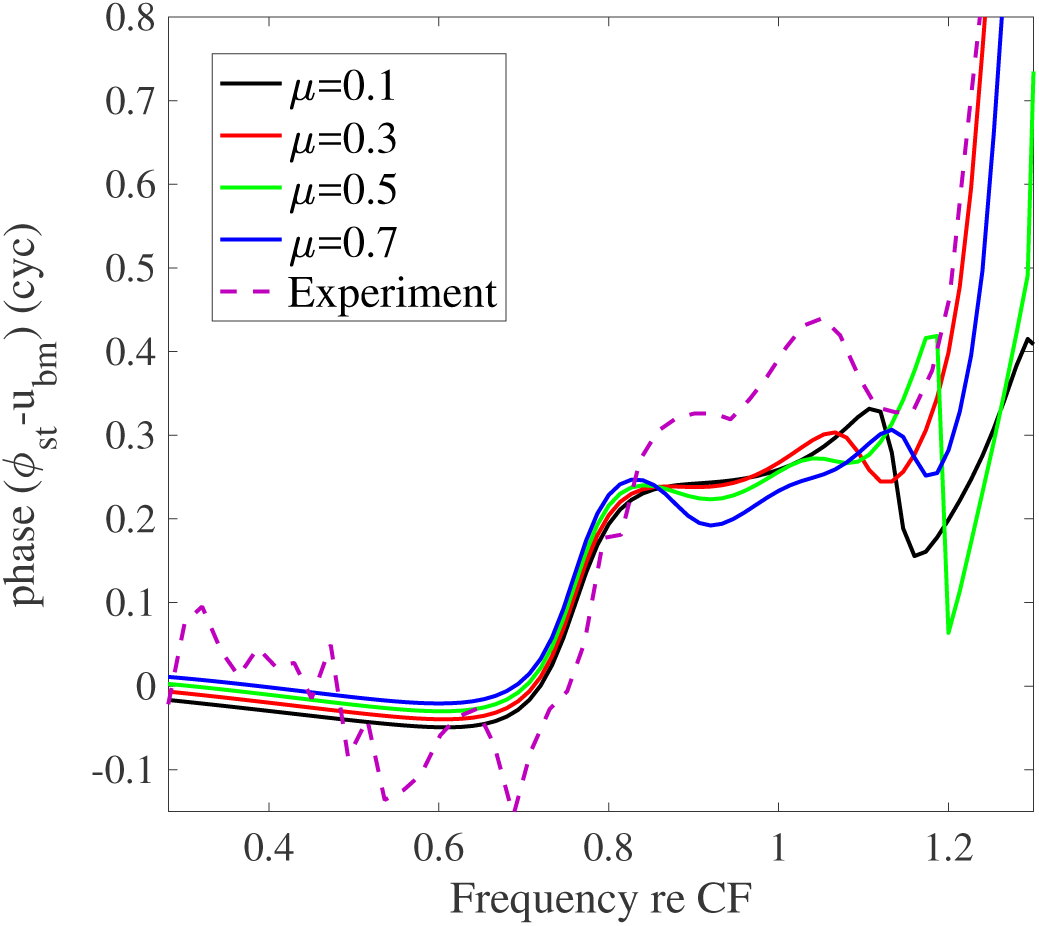
SPL dependence of the phase transition between the ST voltage and BM displacement. The SPL variation in the model is simulated by changing the MET scaling factor *µ*. The experimental value for 60 dB SPL (from (5)) superimposed as a dashed line.

## DISCUSSION

### TM-radial dynamics initiates the cochlear amplifier in the mathematical model

The notch in the amplitude of the voltage response signals both a change in the voltage phase relative to the BM motion and the onset of nonlinearity in the amplitude of the BM displacement. In our mathematical model, the notch in *ϕ*_*st*_ corresponds to a notch in the HB rotation relative to the RL (quantified by *u*_*hb*_, see Fig. 3). According to Eq. 1 in the Methods section, the MET current will follow *u*_*hb*_. The ST voltage (*ϕ*_*st*_ = LCM) is mainly due to the local MET current flowing through the resistance of the cochlear fluids, and, like *u*_*hb*_, *ϕ*_*st*_ also displays a minimum. Both the predictions and experiments show an SPL-independent frequency where the phase shift occurs, suggesting that a passive mechanism is responsible for this feature. In the mathematical model, we are able to explicitly identify this mechanism. The shift frequency occurs at the radial resonance frequency of the TM mass attached only at the limbus (and *not* at the OHC-HB). This frequency is given by 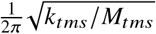 where *k*_*tms*_ is the stiffness of the attachment of the TM to the spiral limbus and *M*_*tms*_ is the TM shear mass (Fig. 3*B*). We determined causality in the model in two ways. First, we directly computed the uncoupled resonance frequency and found it was equal to 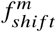. Second, we manipulated the limbal attachment stiffness in the three-dimensional finite element model and were able to predictably adjust the notch frequency according to the resonance calculation above.

To make this explanation more concrete, a simple two-degree-of-freedom abstract model is considered in Fig. 8. Two masses (*m*_1_ and *m*_2_) are connected through elastic and dissipative mechanical elements (springs and dampers). This system represents a simple conceptual model for mechanical interactions between BM, TM and HB as illustrated in Fig. 8*A*, and the parameter values are chosen for demonstration and are not physiological. The mass, *m*_1_, in analogy to the mass of the TM, is anchored to the foundation via *k*_1_ in Fig. 8*B*. The HB deflection in this model is analogous to the difference between *x*_1_ and *x*_2_. The simplicity of this model allows the analytical closed form relation of the mass displacements to be calculated as presented in Eq. 3 (here damping terms are neglected for the sake of simplicity). The denominators of *x*_1_ and *x*_2_ in Eq. 3 are identical and the corresponding roots determine the coupled resonance frequencies of the system. Calculating the relative displacement between the two masses (*x*_1_-*x*_2_) we can see that the numerator is given by *F*_0_(*m*_1_*ω*^2^ − *k*_1_) a term which exhibits a minimum at the frequency which coincides with the resonance frequency of mass *m*_1_ when detached from *m*_2_, 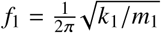.

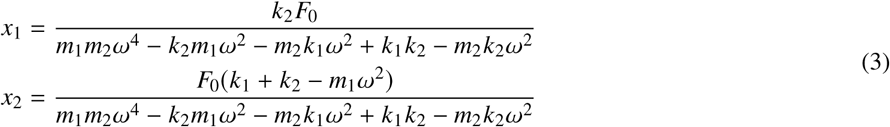

**Figure 8:**
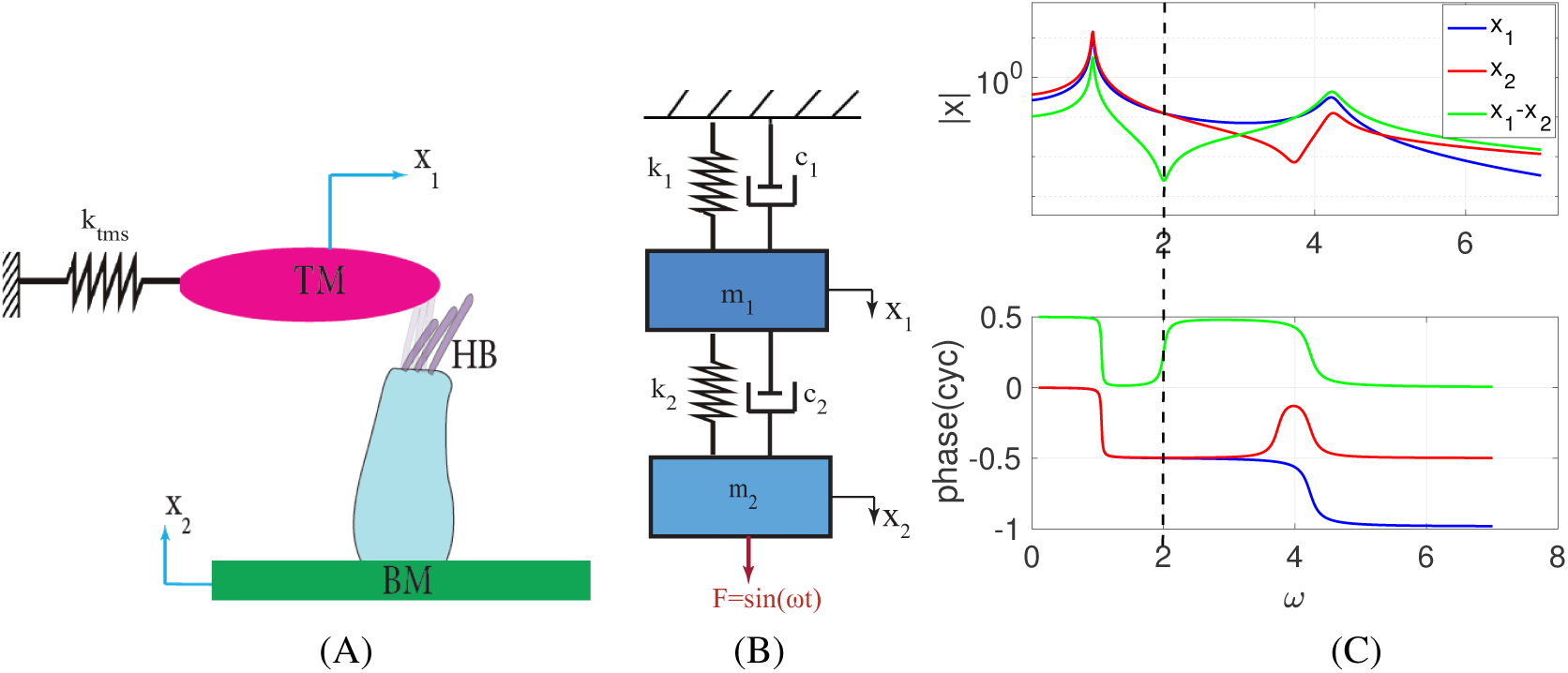
An anti-resonance node in the frequency response of a two degree-of-freedom system describes the notch observed in the complex microstructure of the OCC. (A) schematic of the simplified BM and TM system (B) system schematic composed of two connected masses (m), stiffnesses (k) and dampings (c). (C) frequency responses of the system shown in (B) for parameters: *m*_1_=1, *m*_2_=2, *k*_1_=4, *k*_2_=7, *c*_1_=0.1, *c*_2_=0.1

Figure 8*C* shows the frequency responses of *x*_1_ and *x*_2_ displacements with the values of the mechanical elements denoted in the caption. The responses show two peaks corresponding to the coupled resonances of this simple two-mass system. The quantity *x*_1_-*x*_2_ shows an anti-resonance node that matches the uncoupled resonance of the mass 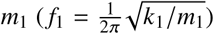. The physical reason for this is that when the unanchored mass *m*_2_ (BM) is forced at frequency *f*_1_, it does not feel any load from *m*_1_ and thus the coupling spring between the masses is undeformed, and the two masses vibrate with similar amplitude and phase. As forcing frequency increases beyond the resonance of the uncoupled TM, the phase of the displacement difference (and, by analogy, the phase of the HB rotation relative to the RL in the cochlear system) undergoes a 180 degree shift because below this frequency the BM amplitude is greater than the TM, and above it the TM amplitude is greater than the BM, and they are moving with a similar phase. The uncoupled resonance of the foundation-supported mass *m*_1_ fixes the frequency where this crossover occurs, independent of the values of the coupling stiffness, *k*_2_, and unanchored mass, *m*_2_. Though the OC is more complicated than the two degree-of-freedom system, the same physical principle applies, in that passive mechanical features (the attachment stiffness and radial TM mass) fixes the notch frequency. The simple model also underscores the fact that significant relative motions (*x*_1_ − *x*_2_) with dramatic frequency dependence occur when the individual motions *x*_1_ and *x*_2_ are not changing dramatically.

Returning to the full cochlear model, the level of damping arising between the shear motion of the RL and TM plays a subtle but key role in shaping the magnitude and phase of the frequency response. In Fig. 9*A*, we investigate the effect on *ϕ*_*st*_ of varying the damping of the subtectorial space. This damping is controlled by *C*_*tms*_, the shear damping of the TM (see Table 2). Increasing the shear-damping factor leads to the elimination of the notch in the magnitude spectrum and reduction of the slope of the overall phase change, but not the amount of the cumulative phase change (Fig. 9*B*). In Fig. 9*B*, we plot the difference between the phase of *ϕ*_*st*_ and *u*_*bm*_ as a function of frequency for different damping values. While the details of the phase change differ, predictions from all three levels show an overall ∼0.25 cycle phase increase in the transition from the notch frequency to CF. Because of this sensitivity to damping, the model predicts animal-to-animal as well as species-to-species variability of the depth (or even existence) of the notch in the magnitude spectrum but not of the overall phase change, which is predicted to be present even in the face of these damping-dependent differences.

**Figure 9:**
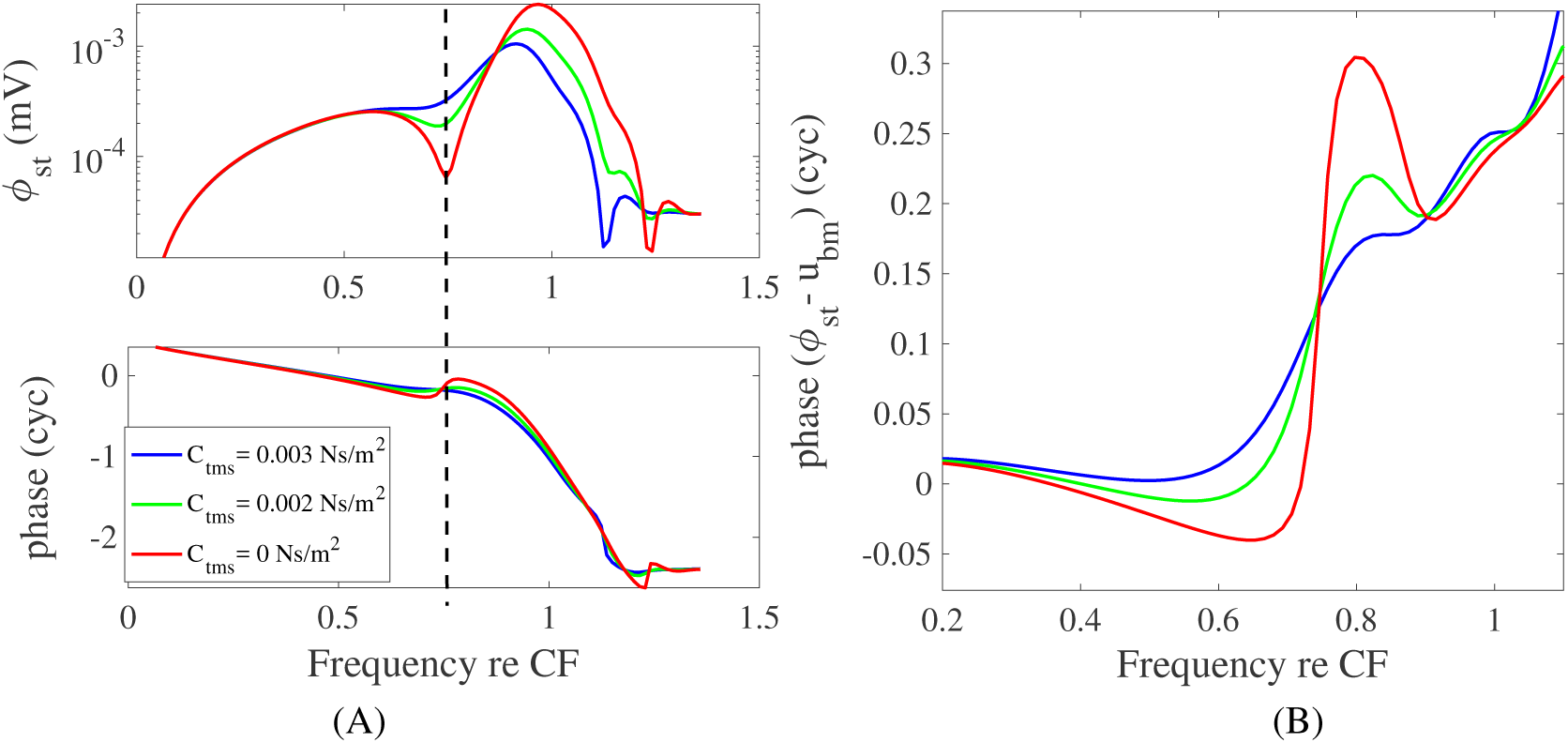
The effect of altering the shear damping between the TM and RL is studied. (A) The frequency dependence of the amplitude and phase of the ST voltage is plotted for varying shear damping in the cochlear model. Increasing the TM shear damping factor (*C*_*tms*_), decreases the depth of the notch (shown by the dashed line) of the ST voltage. (B) The relative phase between the ST voltage and the BM displacement. Increasing shear damping decreases the transitional slope but not overall phase change. (CF = 15.3 kHz)

We used the numerical model to compute the mechanical power that the OHC somatic electromotile force injects (amplifies) or removes (dissipates) at the BM. The energy converting electromotile force is proportional to the cells’ transmembrane potential (40) as described by Eq. 2. Power depends on the relative phase between the force and the motion as well as the magnitude of each. Figure 10 shows the model prediction of this power (Π_*ohc*_) for high-level sound input (using *µ* = 0.1) and low-level sound input (using *µ* = 0.5). A positive value for Π_*ohc*_ indicates power injection (green) from the OHC electromotile force to the BM while negative Π_*ohc*_ represents dissipation (red). The vertical dashed line at 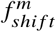 in Fig. 10 is seen to define the boundary between dissipation and amplification. Fig. 10 shows that the OHC power amplification/dissipation normalized to input power is level dependent and thus nonlinear. The level-dependence is greater above the notch frequency (Fig 5B) where the OHC power is positive. The frequency above which power becomes positive is the same for both low and high SPLs. The injected power peaks before the CF. This is because the phase difference between the somatic force (proportional to the OHC transmembrane potential) and BM displacement reaches almost 0.25 cycle at this frequency. Hence, the active force is nearly in phase with the BM velocity; a condition required for the most effective power injection on the BM motion. In addition, the amplitude of the MET current (driven by *u*_*hb*_) is increasing from its local minimum at *f*_*shift*_ and the BM displacement amplitude is also increasing, achieving its maximum at CF. Therefore, the OHCs at a more basal region amplify the CF response of a more apical location, boosting the wave as it passes. Finally, our model predicts above-CF dissipation (the large red region) for low-level sound, indicating the possibility that the active process helps to stabilize the system through dissipation at higher frequencies.

**Figure 10:**
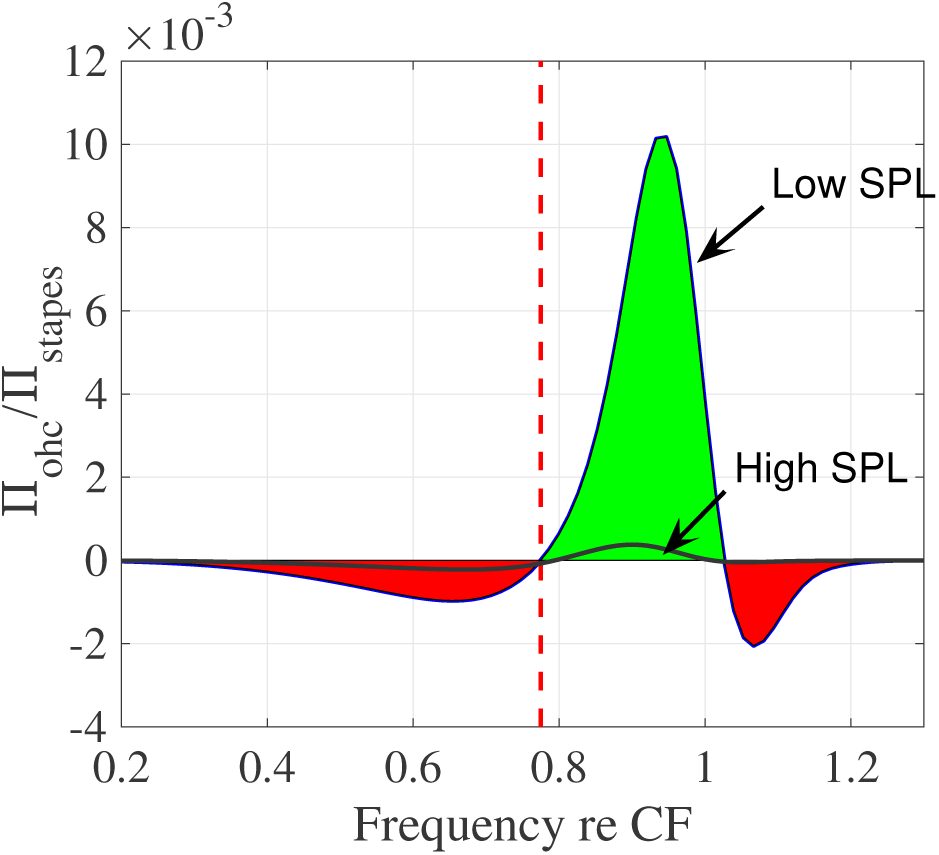
Model prediction of the power exchange between a single OHC and the BM for different activity levels as controlled by the MET scaling factor, *µ*, which is set to 0.1 (for high SPL simulation) and 0.5 (for low SPL simulation). The OHC active power deposited on the BM is calculated as 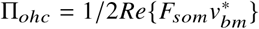, where ^∗^ represents the complex conjugate and *v*_*bm*_ is the velocity of the BM. The OHC power is normalized to the stapes input power. Negative power indicates power dissipation while positive values denote power generative region. The OHC active force is introduced in Eq. 2. The vertical dashed red line is drawn at 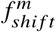 which corresponds to the frequency boundary where the power deposition changes from dissipative (red) to generative (green). Frequencies are normalized to the CF=15 kHz and powers are normalized to the stapes input power (Π_*stapes*_).

### Relation to previous experimental studies

There is one piece of direct *in vivo* evidence that the uncoupled TM radial resonance is roughly ½ octave below the CF of the location along the cochlear spiral. In Lee *et al.*, (6) measurements were made in a living *Tecta*^*C*1509*G*/*C*1509*G*^ mutant mouse (whose TM is detached from the OHC HBs but attached to the limbus). As discussed in the text of that paper regarding Figs. 9*I,J*, there was a “dramatic” shift from phasic to antiphasic motion of the radial motion of the TM relative to the transverse motion of the BM. This is most clearly seen in the movies in the lower two panels of Movie 1 from Lee *et al.* (6), where the radial motion of the TM can be seen to switch from being in phase with the BM motion at 5 kHz to being out of phase at 8 kHz, indicating the passage through a resonance. Since the CF of their measurements was 10 kHz, the hypothesis put forth in the present paper would predict the radial resonance to occur ½ octave below this frequency, at ∼7kHz, which is consistent with the paper’s experimental result. While the experimental results of that study (6) did not show a robust notch in the HB rotation amplitude spectrum of the wild type animal, a clear 0.3-0.5 cycle phase change was seen as the stimulus frequency increased from ½ octave below CF to CF (their Fig. 5*F*). This phase transition was observed in both active and passive preparations, which is consistent with our hypothesis that a passive mechanism underlies this process.

There is substantial indirect evidence of a notch in electrical and neural responses of the cochlea. Ref. (41) discussed several studies that showed a below-CF notch in auditory afferent nerve fiber (ANF) data. ANF measurements in cat (42, 43), mouse (44) and chinchilla (45) all showed a local increase in the threshold (corresponding to a notch in sensitivity) at a frequency approximately a half octave below the CF of the fiber. Measurements of the hair cell voltage responses also have shown a notch below the CF. For instance, the OHC/IHC voltages measured in Fig. 9 of Ref. (46) show a null near 10 kHz for a cell with CF of 17kHz. In another paper from the same group (47) (their Fig. 4), a peak in the threshold pressure needed to attain a given OHC voltage is seen at a frequency of 0.6CF.

In studies in guinea pig in which voltage was measured at the BM and within the OC following measurements of BM motion in the same animal (16), the voltage slightly lagged BM displacement (taking positive displacement towards BM from ST, as in (5)). The voltage responses did not undergo a phase transition at the frequency where BM responses became nonlinear, although at supra-BF frequencies large phase differences were observed, similar to Fig. 7. The difference in the sub-CF results presented in the current work compared to (16) might be related to the fact that the two studies used different animals (gerbil versus guinea pig), electrodes (wide-frequency-band metal versus low-pass-frequency glass), and measurement protocols (simultaneous versus sequential). It will be informative to perform further experiments in guinea pigs and other animals, to specifically explore if this important phase transition that exists in gerbils is present in other mammals.

### Relation to previous models and concepts

Theories emphasizing the importance of the TM in cochlear mechanics have a long history. In particular, Zwislocki and Kletsky (18) theorized that coupling the TM to the BM via a nonlinear spring could produce the sharpness and nonlinearity that could not be achieved with BM mechanics alone. Our modeling results are consistent with predictions from the model of Zwislocki and colleagues (48). In that study, the relative contribution of the TM to the RL motion was manipulated to create notches in the response (Fig. 17 of that paper shows a notch along with a phase jump at a frequency below the CF). However, the OHC active process was not included, and the significance of the notch and the phase jump was not discussed with respect to cochlear amplification. Also in 1980, Allen (19) noted that the zero of a transfer function (which is fixed by the radial resonance of the TM and its limbal attachment) would explain the 180° phase shift observed in some neural data. The work of Allen (19) and Zwislocki (18) predated the discovery of OHC electromotility (49). Hence, their work did not address the question of OHC electromotile force generation and power transfer, a key difference between these models and our model.

An early transmission line model of active cochlear mechanics (20) used two coupled masses to model the OCC and predicted responses that contained a significant sub-CF notch and concurrent phase ripple/shift. The notch and shift occurred in both BM displacement and in hair bundle displacement, and were not present in passive mechanical responses, so are qualitatively dissimilar to our experimental and modeling results, but share basic behavior with our results. Ref. (20) was one of the first models to include OHC electromotility and discuss power gain.

The TM-mechanics-induced phase-shifting mechanism we describe in this paper is in line with analysis of experimental results from Gummer et al., (50), which noted that a notch frequency coincides with the transition of the shear load from the TM onto the OHC HBs from spring-like to inertia-like, with an attendant phase shift that they argued was conducive for amplification. Ref. (41) applied a two-degree of freedom model to analyze experimental data and concluded that the TM vibratory phase and resonance was critical for the timing of the active process. Dong and Olson (5) identified the notch in the ST voltage and analyzed the phase of this quantity in view of the electromotile process and showed that the phase shift activated power amplification by OHCs. In this paper, we use a full cochlear model to link the specific micromechanics to this observation providing the modeling basis for the nonlinear behavior.

### Summary

In this paper, we provide experimental and theoretical evidence supporting the role of the TM as the controlling factor for activation of the cochlear amplifier. Input sound pressure creates a travelling wave along the cochlea which generates vibrations on the OCC components (e.g BM, TM, HB). Deflection of the HBs induces transduction current through the MET channels giving rise to a transmembrane potential across the basolateral membrane of the OHC. The transmembrane potential causes an active somatic force which is then applied to the BM. The effectiveness of this applied force in amplifying the mechanical motions (while taking energy from the electrical domain) relies on the phase relation between the BM and transmembrane voltage of the OHC. We have shown that the passive mechanics of the TM can set the conditions necessary for amplification.

## ACKNOWLEDGEMENTS

The authors thank A. Fridberger for helpful discussions. This work was supported by NIH grants DC-004084 and T32DC-00011 (KG and AN) and R01-DC015362 (EO).

## AUTHOR CONTRIBUTIONS STATEMENT

E.O, A.N, C.E.S. and Y.W conducted the experiments. A.N and K.G developed the models and analyzed the data. E.O and K.G designed the research and supervised the project. A.N. applied mathematical tools, performed the research and drafted the paper. All authors participated in writing and approved the final manuscript.

## SUPPLEMENTAL INFORMATION

### Exploration of possibility that phase cancellation contributed to sub-CF notch and phase shift

We wanted to consider the possibility that the sub-CF notch might be due to phase cancellation of currents from local and distant cochlear locations, rather than a local electromechanical effect. We have explored this possibility in two ways. Our first approach was to exercise our full numerical model, where we can turn off longitudinal coupling of the electrical current (by artificially increasing the resistance in the scalae). In our model, the electrical response is computed using multiple cables (ST, SV, SM) coupled to the active cochlear model (see Fig.S1A and Ref. (33) for the detail of the cable model). In the case when there is no current spread, propagation of current beyond each cross section is not possible and phase cancellation is likewise not possible. With a current spread of zero the notch in the magnitude and the phase change persists (see Fig. S1B-C). When current spread is turned back on (using the normal model parameters) the notch is slightly diminished in depth compared to the “no cables” model. A notch appears above the CF where the phase is varying rapidly, showing that phase cancellation can occur in the model. Therefore, theory predicts current spread is not the main factor for the sub-CF notch and associated phase change.

Second, as in (5), we used a cable model (that paper’s Fig. 7 and 8) to explore the effect of current spread. This approach was also used in Ref. (16) and several other works (51, 52). In the cable model OHC transducer current was assumed to be linearly proportional to BM displacement (a simplifying assumption). Using a post-processing technique, the frequency response of the BM displacement (transducer current) at a single location is transformed to a spatial dependence according to a scale invariant approximation (e.g., following Olson, et al. (53)). The transducer current is then spatially convolved with the current point spread function (Green’s function) with a space constant (80 and 200 microns were explored) to obtain the spatial dependence of the voltage field. We assume a space constant larger than the 42 *µ*m value indicated by (16) for the space constant within the OCC; the larger value is a conservative approximation as it would induce greater phase cancellation. The process and result is in Fig. S2. No sub-CF notches were introduced in the voltage. Based on the results of Figs. S1 and S2, phase cancellation due to current spread does not produce the sub-CF phase shift and amplitude notch in voltage.

**Figure S1:**
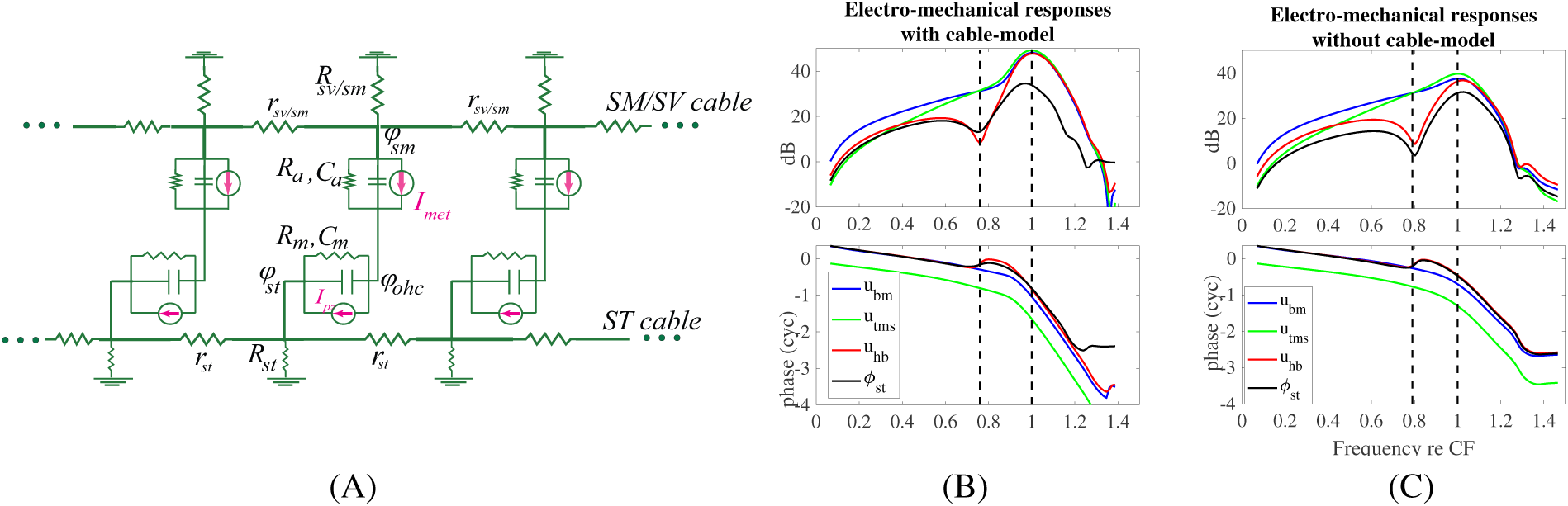
Using the full cochlear model of this paper to examine the possibility that the sub-CF phase shift of voltage relative to BM displacement, and often-accompanying amplitude notch, are due to phase cancellation from distant sources. (A) Electrical network of the OHCs (see Fig 3C) coupled along the cochlea using different cable models on the SM/SV and ST. (B) The full model responses with the cable model. (C) The model responses without the cable model (i.e, no current spread along the cochlea was allowed). The phase shift and voltage notch are present in both cases, indicating they are not due to phase cancellation from interacting electric fields.

**Figure S2:**
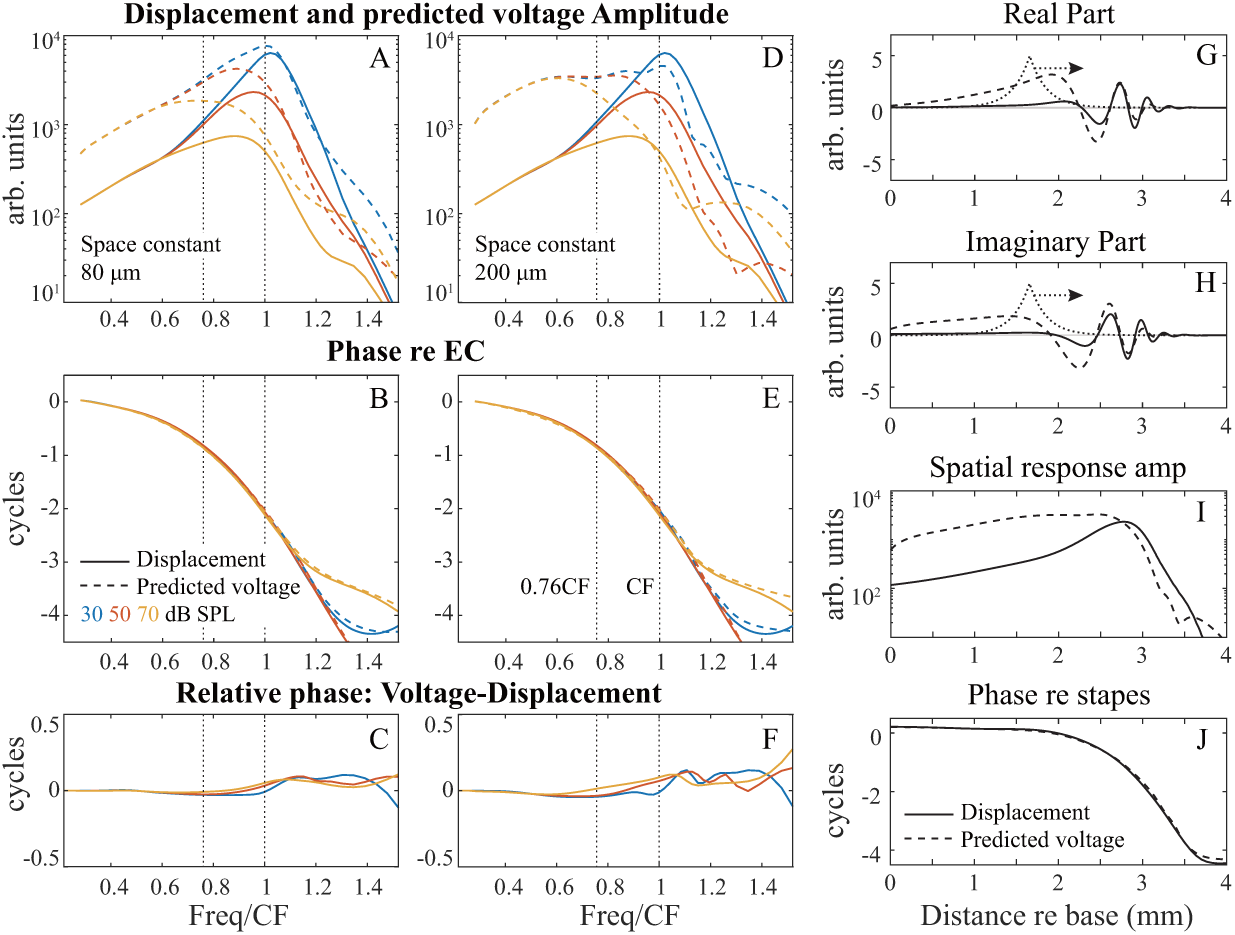
Using the experimental data to explore the potential effect of current spread on the sub-CF notch and phase shift. Two different exponential decay lengths are used, 80 *µm* and 200 *µm*. 80 *µm* was used in Dong & Olson (5) to fit their data, and 200 *µm* is included for illustration because it allows for a larger contribution from distant sources. In panels A-F the color code represents the SPL applied (see panel B) while the displacement field is given by solid lines and the predicted voltage is shown using dashed lines in A,B,D,E. The horizontal axis in A-F is frequency/CF. In G-H the point spread function is shown using a dotted curve and the horizontal arrow represents the convolution operator with the displacement field (solid line). The result of the convolution operation is the predicted voltage field shown using a dashed line. I-J The amplitude and phase are shown for one frequency, versus longitudinal location. The location data are cast back into the frequency domain for the voltage predictions in A-F. Phase cancellation produces supra-CF amplitude notches, but not sub-CF notches. Their location varies with frequency and amplitude. A sub-CF phase shift is not produced.

